# ODDB: Ocular Disease Database for integrated analysis of ocular disease–gene–drug relationships

**DOI:** 10.1101/2025.10.28.685137

**Authors:** Umair Seemab, Anna Kalatanova, Saad Hassan, Ziaurrehman Tanoli, Henri Leinonen

## Abstract

Ocular diseases such as age-related macular degeneration, glaucoma, diabetic retinopathy, and inherited retinal dystrophies are leading causes of vision loss worldwide, yet existing databases often address only limited aspects of these disorders. To fill this gap, we developed the Ocular Disease Database (ODDB), a web-based resource that integrates genes, biomarkers, variants, and drugs associated to ocular diseases. Data were systematically collected through literature mining of PubMed-indexed journals, the NCBI Gene Expression Omnibus (GEO), and drug regulatory agency datasets. Multi-omics, experimental, and clinical information were harmonized using standardized integration workflows. The database is organized according to two complementary ontologies: one based on the anatomical site of pathology (cornea, retina, optic nerve) and another on gene inheritance pattern. ODDB currently covers over 170 ocular diseases, more than 1190 genes, 2400+ variants, and 386 drugs, including both approved and investigational compounds. Each record includes detailed annotations of associated genes, variants, therapeutic targets, and mechanisms of action. The platform supports interactive querying and network-based visualization of disease–gene–drug relationships. All data was internally validated for accuracy and are compliant with FAIR principles, ensuring accessibility and interoperability. ODDB (https://www.oculardiseases.fi/) provides a comprehensive and standardized reference for exploring molecular mechanisms and therapeutic opportunities in ocular diseases.

## 1. Introduction

Eye diseases encompass a broad spectrum of conditions that cause visual impairment and even complete blindness [1]. Common eye diseases include cataracts, glaucoma, diabetic retinopathy, and age-related macular degeneration (AMD). Over 2.2 billion people are estimated to be visually impaired, with many cases being preventable but remaining unaddressed [2], [3], [4]. The challenge with eye diseases, particularly cataracts, is more pronounced in low- and middle-income countries, particularly among disadvantaged rural and ethnic minority populations [5], [6]. The burden of eye diseases is expected to increase substantially with population aging across the globe [7], [8].

While the transparency of the eye and modern imaging technologies offer unique opportunities for early diagnosis and targeted intervention, many eye diseases, particularly blinding retinal diseases, remain untreatable and demand a holistic and multi-disciplinary approach for improved care [9], [10], [11], [12]. In recent years, several public resources have been developed to support ocular disease research. These include the EyeDiseases database, which integrates multi-omics data across a wide range of eye conditions [13]. Another resource is the Eye Biomarker Database (EBD), which catalogs disease-associated genes and clinical annotations [14]. Additionally, the Eye Transcriptome Atlas offers tissue and cell-type specific gene expression profiles across various compartments of the human eye [15]. Additionally, broad-spectrum platforms such as Open Targets [16] and DisGeNET [17] provide valuable insights into disease–gene associations across multiple therapeutic areas.

However, these generalized databases lack a dedicated, eye disease–focused resource that comprehensively integrates disease–gene–drug associations under one platform [18]. Such a platform is crucial for bridging the gap between genomic data and translational applications. Here, we have developed an interactive and comprehensive online platform that integrates biomarkers, FDA-approved drugs, clinical trial data, and genetic associations specific to ocular conditions. Unlike existing platforms that primarily focus on either genetic or expression data, our tool emphasizes translational relevance, linking molecular data directly to therapeutic insights. Continuously updated, the platform is designed to facilitate therapeutic discovery, advance patient care in ocular diseases, and provide a robust resource for data-driven ophthalmic research

## 2. Materials and Methods

### 2.1. Classification ontology for ocular diseases

The structural design of our ocular disease database is based on a dual categorization framework that enables comprehensive disease annotation and intuitive data navigation. First, diseases were categorized based on the anatomical location of affected ocular tissues (e.g., retina, lens, cornea, optic nerve) and guided by ocular disease–specific vocabulary drawn from Medical Subject Headings (MeSH) [19] and Online Mendelian Inheritance in Man (OMIM) [20]. This classification enables precise alignment with existing biomedical terminologies and facilitates integration with other disease-related resources. Over 170 curated ocular diseases were classified into 12 major anatomical categories (**Figure 1A**). In parallel, an inheritance-based categorization was implemented by analyzing the mode of inheritance of associated genes, such as autosomal dominant, autosomal recessive, or X-linked as illustrated in **Figure 1B**, allowing users to explore the database from both anatomical and genetic perspectives. This framework facilitates exploration of the genetic etiology of ocular diseases and supports hypothesis generation for further research.

**Figure 1:**
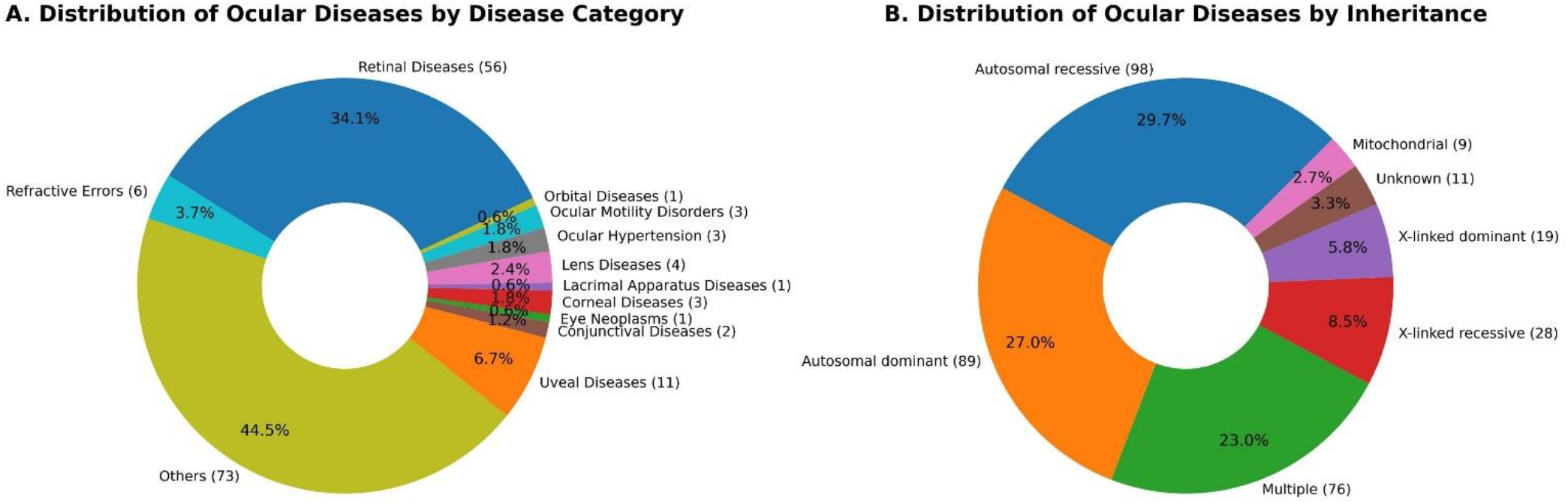
**(A)** Distribution of ocular diseases by disease category. Retinal diseases represent 34.1% of all entries, followed by “Others” (44.5%) and smaller categories like uveal, corneal, and lens disorders. The “Others” group includes syndromic and systemic diseases with ocular involvement. **(B)** Distribution of ocular diseases by inheritance. Autosomal recessive inheritance accounts for 29.7% (98 diseases), followed by autosomal dominant (27.0%, 89 diseases) and multiple inheritance modes (23.0%, 76 diseases). X-linked recessive (8.5%) and dominant (5.8%) patterns are less common, while mitochondrial (2.7%) and unknown (3.3%) categories are rare.

### 2.2. Data Preparation and Curation

Eye-related disease information was first collected from OMIM and MeSH databases to establish the primary disease list. Sequencing datasets corresponding to these diseases were retrieved from the Gene Expression Omnibus (GEO) [21] to obtain gene expression and variant-level information. The raw sequencing data were processed using an in-house RNA sequencing pipeline following standard practices, including quality control with FastQC [22], read alignment with STAR aligner [23], feature counting using HTSeq [24]. For variant identification, aligned reads were processed using SAMtools [25] and GATK best-practice workflow [26], which included duplicate marking, base quality recalibration, and variant calling using HaplotypeCaller. The resulting variant call format (VCF) files were filtered and annotated with Ensembl Variant Effect Predictor (VEP) [27] to identify functional and pathogenic variants. Normalization and differential expression analysis with DESeq2 [28]. The processed data was curated and used as a test dataset for model training.

For text-based data extraction, a natural language processing (NLP) approach was adopted. BioBERT [29], a biomedical domain-specific language representation model, was fine-tuned using the curated ocular disease literature to automatically extract disease–gene–variant relationships from PubMed Central [30] and genome-wide association data from the GWAS Catalog [31]. Drug information associated with identified diseases and genes was gathered from ChEMBL [32] and ClinVar [33], following similar extraction procedures as used in Repurpose Drugs [34].

All data sources were standardized and manually verified by annotators to ensure consistency and accuracy of labels. The manually curated and standardized data served as the training set for the BioBERT-based NLP classifier. The model was iteratively trained and refined by comparing automated predictions with expert annotations, and the final validated dataset was generated through this human-in-the-loop approach (**Figure 2**).

**Figure 2:**
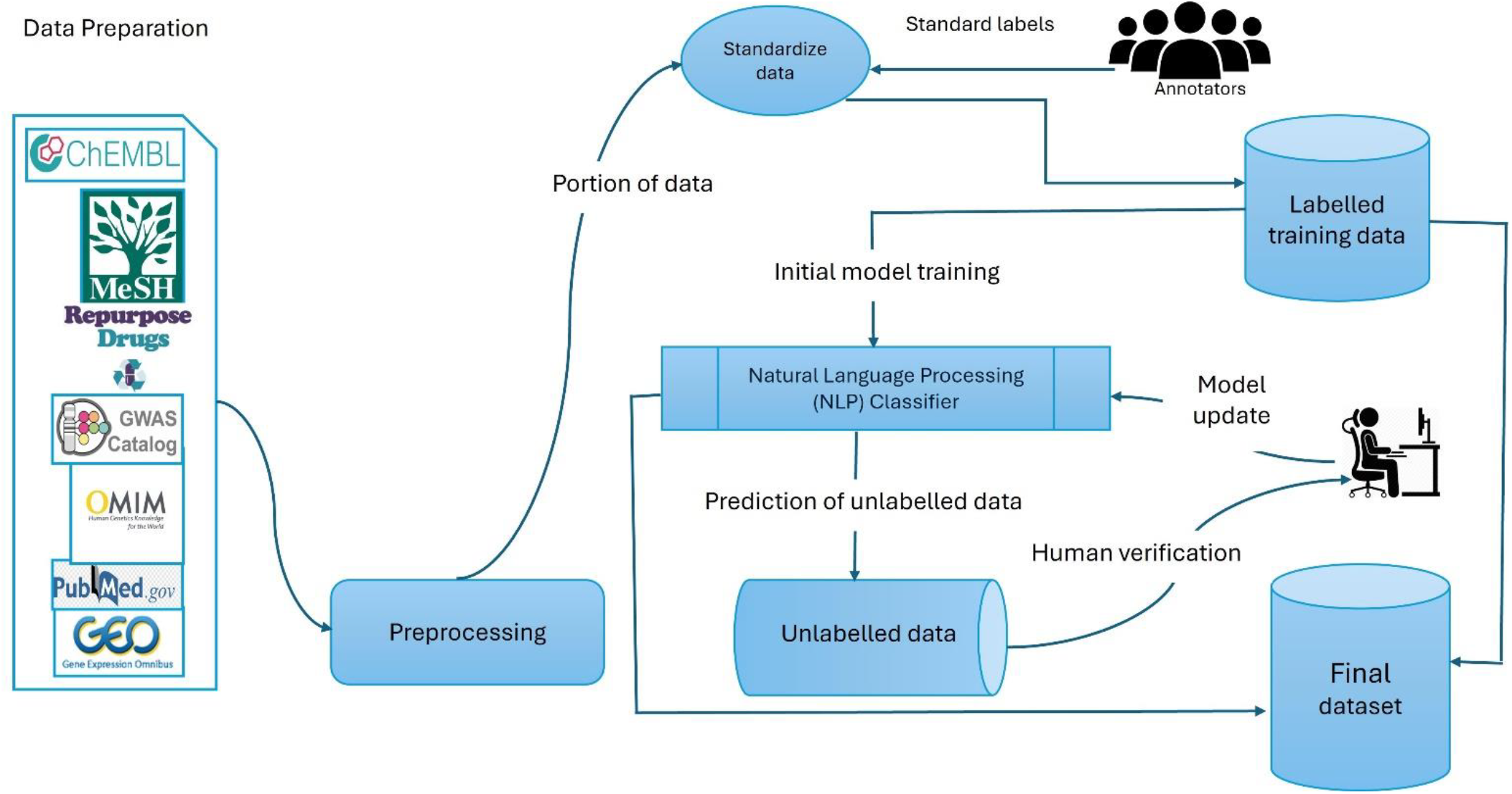
Data collection and annotation workflow.

### 2.3. Web platform development

The front-end was built using React (v18), offering a dynamic and responsive interface that adapts seamlessly across devices. For network visualization, we employed the powerful D3.js (v7) library, which allowed us to create rich, interactive graphs to effectively represent complex ocular data. To maintain a consistent and responsive design across different screen sizes, we integrated Bootstrap (v5.3.5).

For efficient data handling, structuring, and transformation, we utilized JSON-based techniques, enabling smooth communication and integration between various components of the application. Instead of using a traditional database engine, we opted for a simpler approach by utilizing CSV files, as the data structure is relatively straightforward and does not require a complex storage solution.

## 3. Results

### 3.1. Overview of ODDB

The Ocular Diseases Database (ODDB) is an interactive web-based platform designed to integrate and visualize data related to ocular diseases, genes, variants, and drugs. The database is structured into six main modules: Home, Analysis, Statistics, Tutorial, Download, and Contact (**Figure 3**).

**Figure 3:**
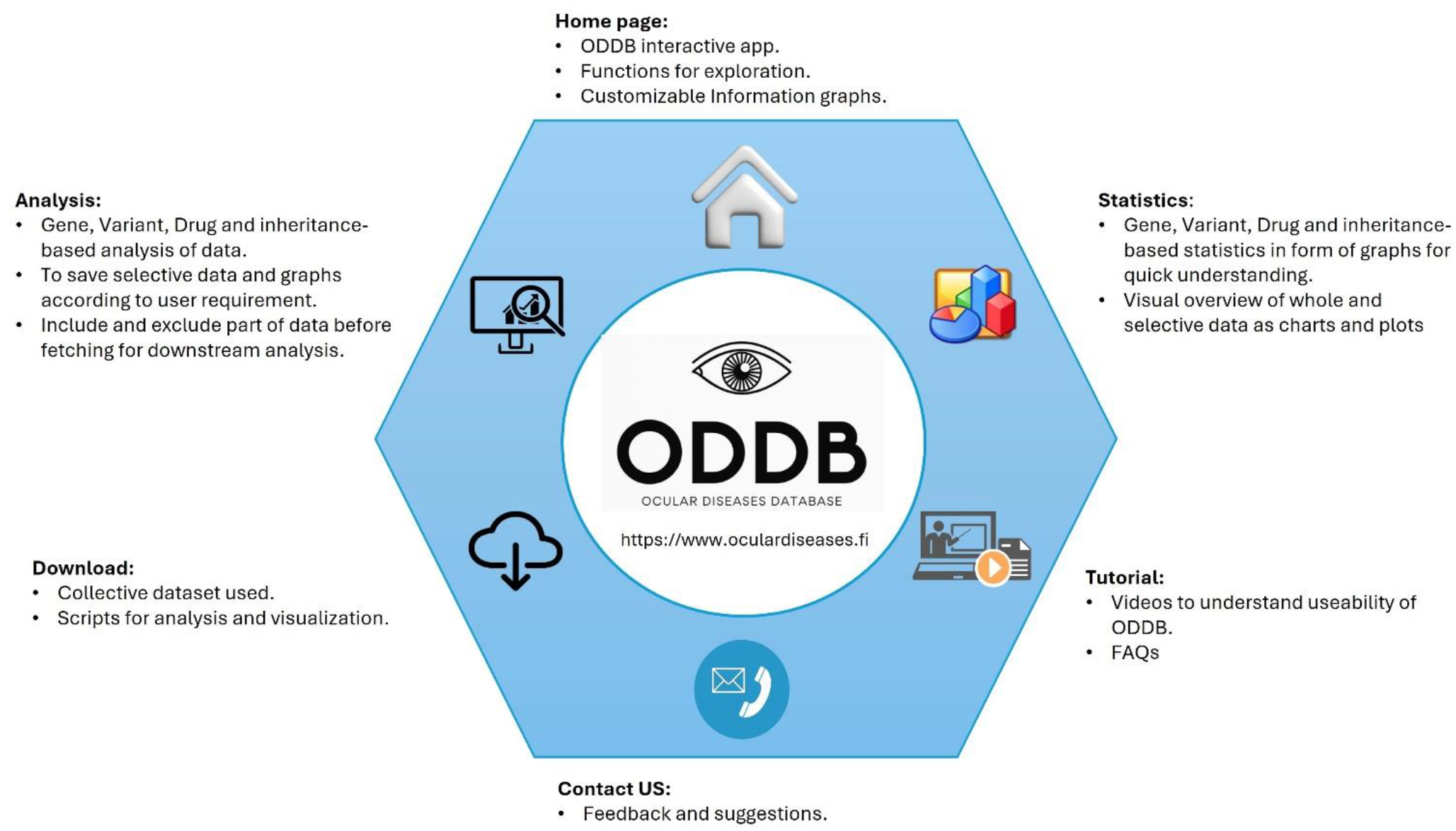
Interface with the ODDB webpage.

The Home module provides access to the ODDB interactive application with customizable information graphs and user-friendly exploration tools. The Analysis module allows users to perform gene, variant, drug, and inheritance-based analyses. It supports selective data saving, inclusion and exclusion of data subsets, and preparation for downstream analysis according to user requirements. The Statistics module offers visual summaries of gene, variant, drug, and inheritance data through dynamic charts and plots for quick understanding. The Tutorial module contains video guides and FAQs to help users understand and navigate ODDB functionalities effectively. The Download module provides access to collective datasets and associated analysis or visualization scripts. The Contact module enables users to share feedback and suggestions for database improvement.

### 3.2. Statistical Overview of ODDB Dataset

The curated ODDB dataset integrates genomic, clinical, and pharmacological data to provide a comprehensive overview of ocular diseases (Supplementary Data files are provided for detailed reference). The statistical analyses illustrate the chromosomal, inheritance, and therapeutic landscape of 170 curated ocular disorders and their associated genes, variants, and drugs.

Overall, ODDB captures more than 1190 unique genes and over 2400 variants mapped across chromosomes 1–22 and X, linked to over 170 ocular diseases which are illustrated in **Figure 4**.

**Figure 4:**
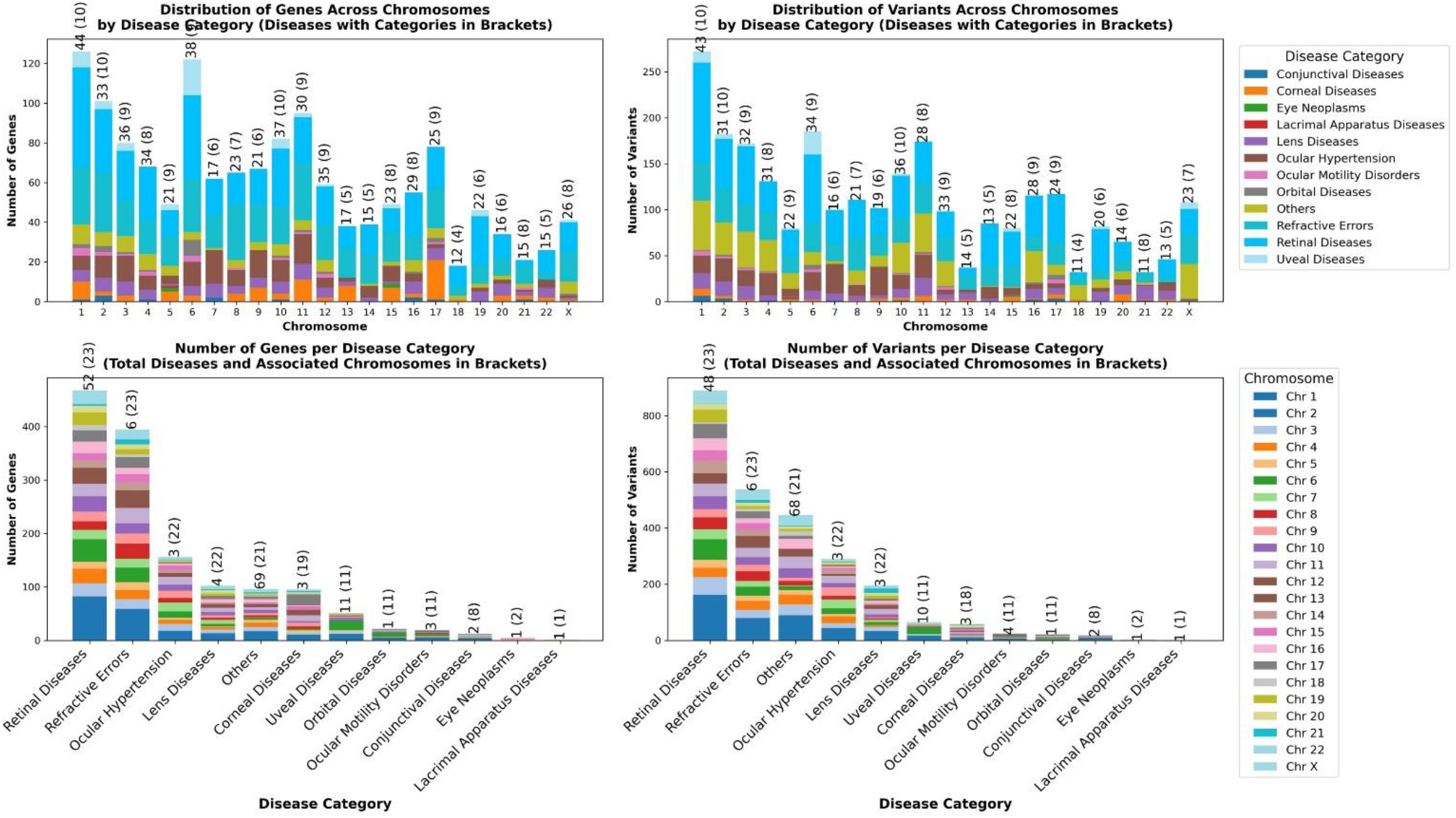
Comparative distribution of genes and variants across chromosomes and disease categories in ocular diseases. **(A)** Distribution of genes across individual chromosomes, stacked by ocular disease categories. Numbers above the bars indicate the total number of diseases and distinct disease categories associated with each chromosome. **(B)** Distribution of variants across individual chromosomes, stacked by ocular disease categories, with total disease and category counts displayed above. **(C)** Number of genes identified per disease category, stacked by associated chromosomes. The total number of diseases and chromosomes involved in each category are indicated above the bars. **(D)** Number of variants identified per disease category, stacked by associated chromosomes, with total disease and chromosome counts shown on top. Colors represent either disease categories (for A and B) or chromosomes (for C and D). The legends on the right show the color codes for disease categories (top) and chromosomes (bottom).

A total of 386 drug compounds, spanning preclinical to approved stages, are catalogued. Retinal and refractive categories account for most gene and drug associations, emphasizing their central role in ocular disease genetics and therapeutic development. To examine the distribution of genetic and therapeutic associations in a subset of ocular diseases, we analyzed the top twenty disorders based on the number of associated genes, variants, and drugs. The number of genes and variants per disease was visualized as stacked bar charts by chromosome (Figure 5, panels A–B). Drug-related analyses revealed the top twenty diseases with the highest number of drugs in clinical development and the top twenty drugs tested across multiple ocular diseases (Figure 5, panels C–D). Bars were color-coded by clinical trial phase (0–4), showing that most ocular drug candidates are concentrated in phases 2 and 3, with only a fraction reaching phase 4, and thus, acceptance for clinical use.

**Figure 5:**
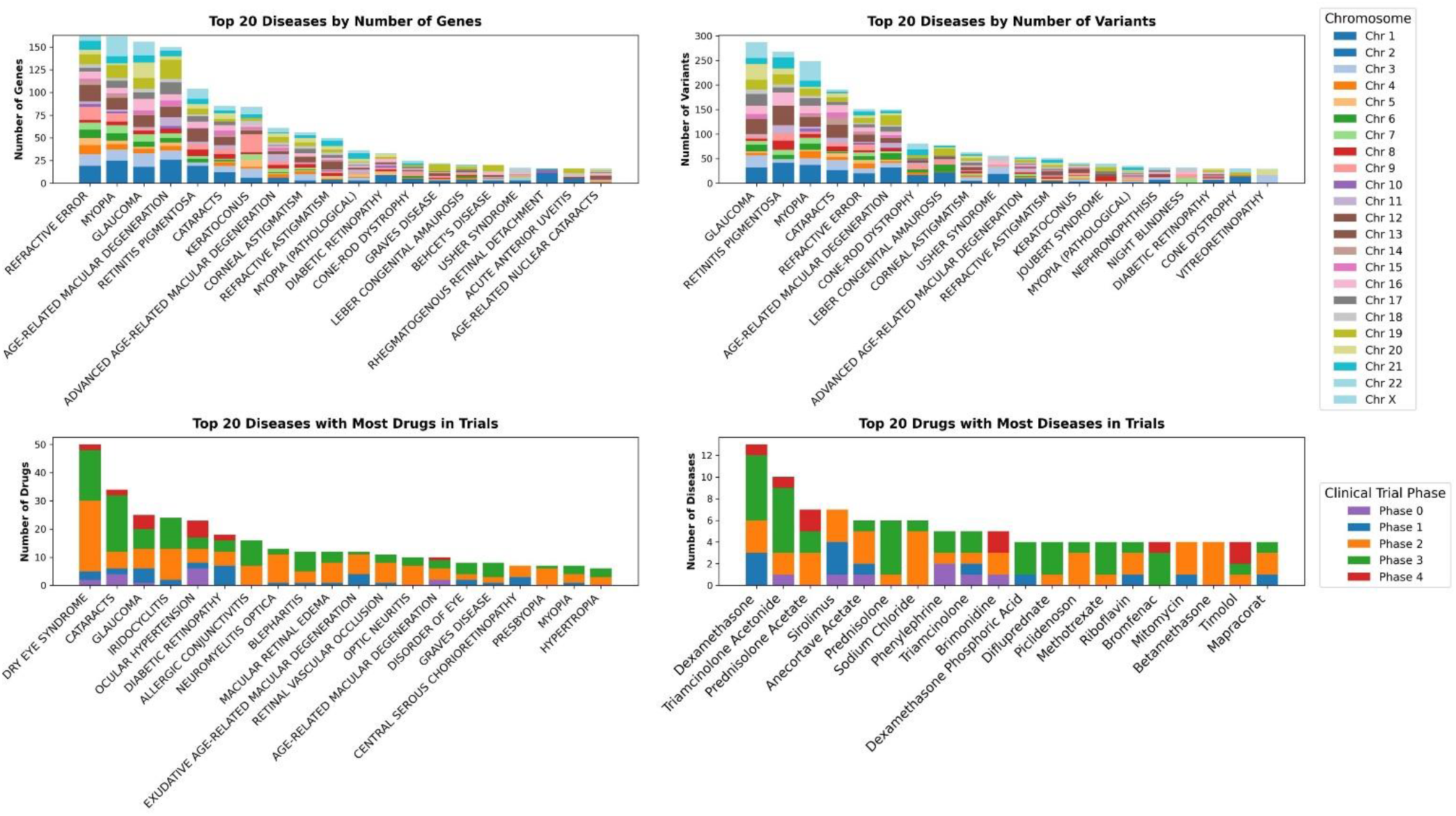
Comparative overview of genes, variants, and drug associations in ocular diseases. **(A)** Top 20 ocular diseases ranked by the number of associated genes, with bars stacked by chromosome. **(B)** Top 20 ocular diseases ranked by the number of associated variants, with bars stacked by chromosome. **(C)** Top 20 ocular diseases with the highest number of drugs in clinical development, with bars stacked by clinical trial phase. **(D)** Top 20 drugs evaluated for the largest number of ocular diseases, with bars stacked by clinical trial phase. Chromosome color codes and clinical trial phase color keys are shown on the right.

## 4. Discussion

The Ocular Disease Database (ODDB) is an open-access web resource that integrates genomic, clinical, and pharmacological data related to ocular diseases. It links diseases, genes, variants, and drugs through an interactive visualization interface to facilitate translational and precision research.

ODDB currently includes 170 ocular diseases, 1,190 genes, over 2,400 variants, and 386 drugs, systematically curated from public resources such as OMIM, DisGeNET, Open Targets, DrugBank, and ChEMBL. Diseases are organized by anatomical localization, including the retina, cornea, and optic nerve, enabling targeted exploration and retrieval. Each disease entry includes curated genes and variants linked to external reference databases and PubMed literature, providing validation and facilitating deeper analysis. These datasets clarify molecular mechanisms, inheritance patterns, and therapeutic targets, strengthening the understanding of genotype–phenotype relationships.

The database also contains detailed pharmacological information on approved and investigational compounds. Each drug entry specifies therapeutic targets, mechanisms of action, and clinical trial phases, allowing users to assess therapeutic strategies and identify potential drug repurposing opportunities. Integration of clinical trial data highlights current research and development trends in ocular therapeutics. Interactive, network-based visualizations distinguish ODDB from static repositories. Network-based visualizations reveal shared pathways, cross-disease gene associations, and drug-target overlaps, enabling intuitive interpretation of complex biological relationships.

A standardized curation workflow ensures data quality, consistency, and scalability. Manual and automated pipelines, including BioBERT-based text mining and cross-validation with multiple databases, were used for integration. The resulting datasets are freely downloadable in structured formats, supporting computational modeling and downstream analyses. ODDB follows FAIR principles findable, accessible, interoperable, and reusable, and is maintained as an open-source resource, ensuring transparency, external integration, and community-driven updates.

The Supplementary Data files provide comprehensive reference material, including gene– disease (Supplementary Data 1), variant–disease (Supplementary Data 2), inheritance-based classification (Supplementary Data 3), drug-disease (Supplementary Data 4), and disease reference lists (Supplementary Data 5). These files enhance reproducibility and support independent validation of the ODDB resource. While ODDB covers major ocular diseases, data on rare disorders and non-coding variants remain limited. Future updates will extend data coverage, include proteomic and metabolomic information, and enhance visualization performance.

ODDB serves as a living platform for ocular disease research. By integrating curated multi-source data, interactive tools, and FAIR-compliant accessibility, it provides a sustainable foundation for discovery, clinical interpretation, and therapeutic innovation in ophthalmology.

## 5. Data availability

www.oculardiseases.fi/

